# Potassium Glutamate and Glycine Betaine Induce Self-Assembly of Sliding Clamps into Higher Order Oligomers

**DOI:** 10.1101/2020.03.05.975235

**Authors:** Anirban Purohit, Lauren G. Douma, Linda B. Bloom, Marcia Levitus

## Abstract

Sliding clamps are oligomeric ring-shaped proteins that increase the efficiency of DNA replication. The stability of the *Escherichia coli* β-clamp, a homodimer, is particularly remarkable. The dissociation equilibrium constant of β is of the order of 10 pM in buffers of moderate ionic strength. Coulombic electrostatic interactions have been shown to contribute to this remarkable stability. Increasing NaCl concentration in the assay buffer results in decreased dimer stability and faster subunit dissociation kinetics in a way consistent with simple charge-screening models. Here, we examine non-Coulombic ionic effects on the oligomerization properties of sliding clamps. Replacing NaCl by KGlu, the primary cytoplasmic salt in *E. coli*, results in the formation of assemblies that involve two or more rings stacked face-to-face. Results can be quantitatively explained on the basis of unfavorable interactions between KGlu and the functional groups on the protein surface, which drive biomolecular processes that bury exposed surface. Similar results were obtained with the *S. cerevisiae* PCNA sliding clamp, suggesting that KGlu effects are not specific to β. Clamp association is also promoted by glycine betaine, a zwitterionic compound that accumulates intracellularly when *E. coli* is exposed to high concentrations of extracellular solute. Possible biological implications are discussed.

## INTRODUCTION

Sliding clamps are oligomeric ring-shaped proteins that increase the efficiency of DNA replication by encircling DNA while tethering DNA polymerases to their templates (1–4). Despite little homology at the primary amino acid sequence level, the structures of sliding clamps are remarkably conserved from bacteria to humans (5–7). The *Escherichia coli* β-clamp (hereafter called β) is a homodimer that contains six globular domains connected by protein loops in two groups of three (Fig. 1A) (7). Consistent with their biological role as processivity factors, sliding clamps are usually stable as rings in solution (8,9). The stability of β is particularly remarkable; 1 nM solutions of β show negligible dissociation over 48 h of incubation at room temperature in buffers of moderate ionic strength, and the dissociation equilibrium constant of the dimer has been estimated at 6.5–65 pM (9,10). We have previously reported the results of a study aimed to determine what factors contribute to the remarkable stability of the clamp interfaces (10). Increasing NaCl concentration in the assay buffer results in decreased dimer stability and faster subunit dissociation kinetics in a way consistent with simple charge-screening models (i.e. Debye-Hückel screening). These and other results led to the conclusion that electrostatic interactions at the dimer interface contribute to the stability of β (10).

**Figure 1:**
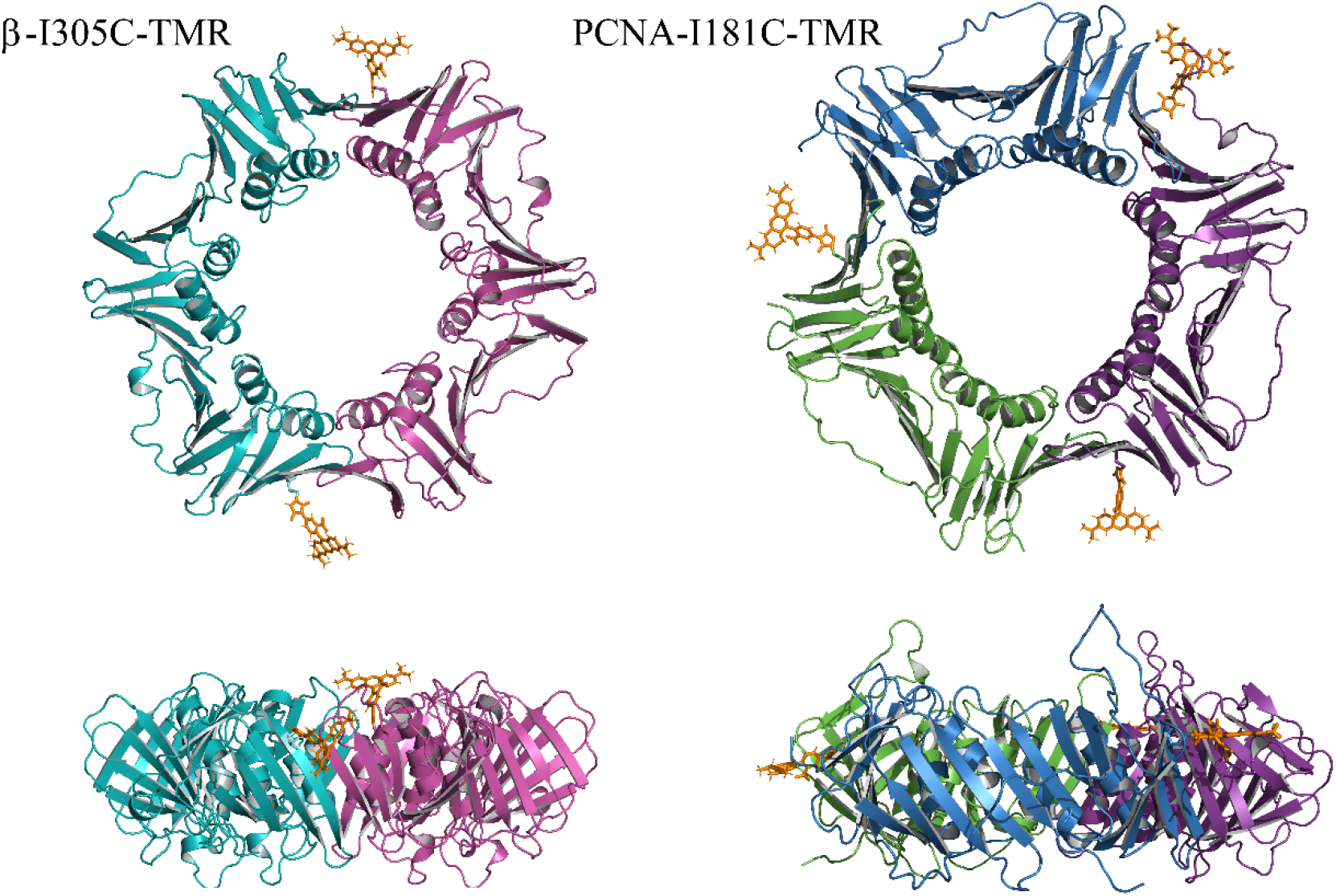
Two perpendicular views of the ribbon diagrams of the two clamps used in this work. Left: *E. coli β*-sliding clamp (PDB: 1MMI). Residue I305 of each subunit was mutated to Cys and a covalent bond between Cys and the maleimide moiety of TMR was created using Pymol. The structure of TMR was obtained from the x-ray crystal structure of tetramethylrhodamine-5-maleimide-actin (PDBeChem id: RHO). Right: *S. cerevisiae* PCNA sliding clamp (PDB: 1SXJ). The crystal structure contains the clamp loader (replication factor C) bound to the sliding clamp, and these subunits were removed from clarity. Residue I181of each subunit was mutated to Cys and covalent bonds between Cys and TMR were created in Pymol.

Ion effects on protein stability have diverse physical origins. At salt concentrations below *c.a*. 100 mM, protein-ion interactions are typically governed by nonspecific electrostatic effects, and their sensitivity to salt concentration can be described in terms of charge screening models. Because Coulombic effects are valence specific but otherwise do not depend on the chemical identity of the ionic species, the same effects are expected for any 1:1 electrolyte such as NaCl. Yet, there are numerous examples in the literature of ion-specific (non-Coulombic) effects on protein solubility, protein stability, and enzymatic activity, that are typically observed around or above *c.a*. 100 mM (11–13). In this article we focus our attention to the potential specific effects of the ion glutamate (Glu^-^) on the stability of β, the *E. coli* processivity clamp. Glutamate is the major low-molecular weight cytoplasmic anion in *E. coli* (14–16), comprising over 40% of the total measured metabolome in exponentially growing cells (14). Glutamate concentration in glucose-fed exponentially growing *E. coli* is about 100 mM (14), but its concentration increases to more than 300 mM in response to osmotic stress (15,17). *In vitro*, KGlu stabilizes folded proteins, protein-protein and protein-nucleic acid complexes (18–26). The accumulation of Glu^-^ and other osmolytes under osmotic stress is believed to have evolved as a mechanism to mitigate the destabilizing effects of high ionic strength on nucleic acid-protein interactions and other interactions among biomolecules (27–29).

Our results show that replacing NaCl by KGlu not only prevents clamp dissociation, but also induces the association of two or more clamps into a protein assembly. Clamp assembly is also induced by glycine betaine (GB), a zwitterionic compound that accumulates intracellularly when *E.coli* and most other living cells are exposed to high concentrations of extracellular solutes (30–32). Results are discussed in the context of a thermodynamic model that takes into account the relative binding affinity of the protein surface for water and for Glu or GB.

## MATERIALS AND METHODS

### Purification of β and PCNA

Genes encoding β and PCNA were mutated using the QuikChange mutagenesis kit (Stratagene) following the manufacturer’s directions to introduce a single surface Cys residue in the proteins to permit site-specific labeling by a maleimide derivative of a fluorophore (33–35). Native Cys residues on the surface β (Cys-260 and Cys-333) and PCNA (Cys-22, Cys-62, and Cys-81) were replaced by Ser. A Cys residue was introduced into β (Ser-109 to Cys or Ile-305 to Cys) and PCNA (Ile-181 to Cys) to incorporate one fluorophore per monomer. These β clamp and PCNA mutants were purified as described previously (35,36)

### Fluorescent labeling of β and PCNA

The engineered Cys residue in β or PCNA mutants was covalently modified with tetramethylrhodamine-5-maleimide or Alexa Fluor 546 C5 maleimide (ThermoFisher) and purified from the free fluorophore as described previously (9,10). Protein concentrations were determined using unlabeled wild-type proteins (β or PCNA) as standards in a Bradford assay (BioRad). Fluorophorophore concentrations were determined from absorbance measurements and molar absorptivities as described previously (9,10).

### Chemicals

The following chemicals were obtained from Sigma-Aldrich (Milwaukee, WI): potassium glutamate monohydrate > 99% pure, mono sodium glutamate salt > 99% pure, betaine monohydrate > 99% pure. Sodium chloride 99% pure obtained from VWR Scientific. tris(hydroxymethyl)aminomethane hydrochloride was obtained from Fisher Scientific.

### FCS Setup and data collection

FCS measurements were carried out using a home-built confocal optical setup. The output of a 532 nm CW laser (Coherent Compass 215M-10, Santa Clara, CA, USA) was expanded, collimated, and directed via a dichroic filter into an Olympus PlanApo 100X/1.4NA Oil objective. The laser power measured before the entrance of the objective was 110 μW. Samples were placed into 50 μL-perfusion chambers pre-treated with BSA to minimize protein adsorption onto the cover glass (see below). Fluorescence was collected via the same objective, separated from reflected excitation light through the dichroic filter, and reflected into a 50 μm pinhole. The emission was then focused into an avalanche photodiode detector (Perkin-Elmer Optoelectronics, SPCM-AQR14). A bandpass filter was used before the detector to minimize background (Omega 3RD560-620). The recorded fluorescence signal was autocorrelated in real time using an ALV5000/EPP USB-25 correlator (ALV GmbH, Germany). The autocorrelation function acquired in this way is defined as: 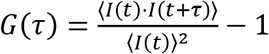, where *I*(*t*) is the intensity at a given time *t* and *I*(*t* + τ) is the intensity measured after a lag time τ. Angular brackets denote averages over the whole measurement. Data was collected for 10 minutes by acquiring twenty 30-second long autocorrelation traces. The 20 autocorrelation decays were averaged prior to analysis as described below. Glass slides (Fisherbrand) were first treated in an ozonator chamber (BOEKEL UV Clean 135500) for 20 min (10 min per side), then sonicated for 45 minutes in 3% Hellmanex (Hellma) cleaning solution, washed extensively with nanopure water and dried under N_2_ gas. Solutions were placed inside silicone perfusion chambers (CoverWell) pressed on top of clean glass slides. Perfusion chambers were first treated with BSA (0.1 mg/mL) for 10 minutes. The chamber was then allowed to dry for an additional 10 minutes after removing the BSA solution, and then filled with the solution of interest for the FCS measurement. Chambers were reused, and rinsed with nano-pure water between measurements until a measurement with water resulted in no correlation and negligible background fluorescence. The confocal parameters were calibrated daily by measuring the FCS decay of a 20 nM TAMRA (carboxylic acid of tetramethyl rhodamine) free dye solution (Molecular Probes, Inc.) with a known diffusion coefficient (D = 420 μm^2^s^−1^) (37). The average decay of the TAMRA solution was fitted using the equation 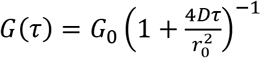 to determine *r_0_*. We note that although the determination of τ_App_ does not require knowledge of *r*_0_, we calibrate the setup daily to ensure *r*_0_ remains constant and thus to be able to compare τ_App_ values measured for independent but otherwise identical experiments.

All reported τ_App_ values are averages of at least three independent experiments. The data of Fig. 2A (β in KGlu) represents the average of 7 independent repeats. Error bars represent standard deviations.

**Figure 2:**
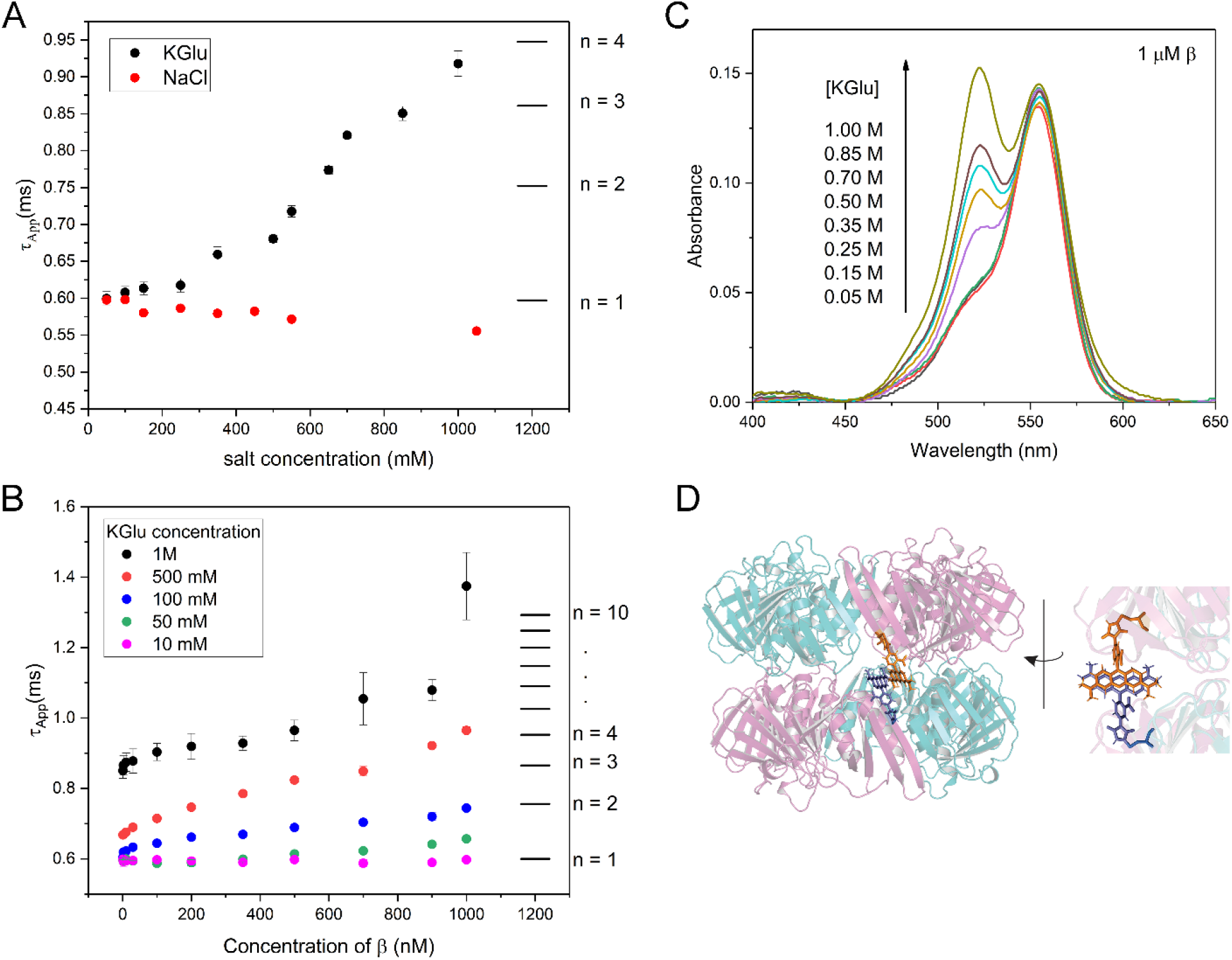
**(A)** Apparent diffusion times (τ_App_) obtained from the FCS decays measured with 1 nM solutions of β-I305C-TMR in an assay buffer containing 20 mM Tris-HCl (pH 7.5) and a variable concentration of NaCl or KGlu in the 50 mM – 1M range. The horizontal lines on the right side of the plot indicate approximate diffusion times expected for protein assemblies containing *n* = 1, 2, … clamps. **(B)** Apparent diffusion times (τ_App_) of solutions containing 1 nM β-I305C-TMR and variable concentrations of unlabeled β. The concentration specified in the abscissa represents the sum of the two. The assay buffer was 20 mM Tris-HCl (pH 7.5) containing variable concentrations KGlu in the 10 mM – 1M range. **(C)** Absorption spectra of 1 μM β-I305C-TMR in TRIS buffer containing 50 mM – 1 M KGlu. **(D)** PyMol model created by placing two clamps on top of each other and rotating single bonds in the maleimide moiety of TMR to obtain a conformation compatible with the strong 520 nm band observed in the UV-Vis spectrum of 1 μM β-I305C-TMR at [KGlu] ≥ 350 mM.

**Fig. 3:**
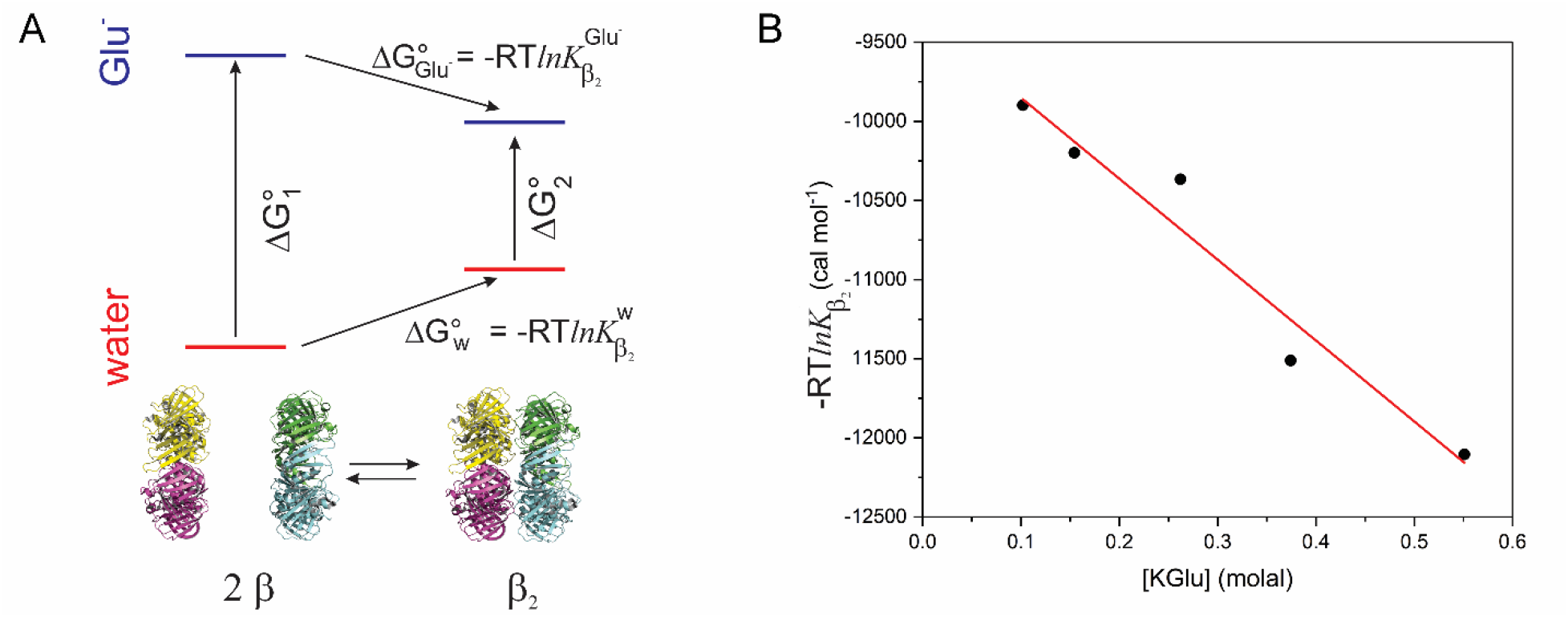
**(A)** Free energy diagram of the effect of KGlu on clamp assembly. Vertical lines represent the standard free energy changes when two *β* clamps 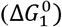 or the dimer 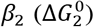 are hypothetically transferred from water to the glutamate solution. 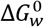 and 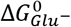 are the standard free energy change for the association of two clamps in water and in the presence of Glu^-^, respectively. 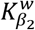 and 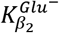 are the thermodynamic equilibrium constants for the formation of the dimer: 2*β* ⇌ *β*_2_ **(B)** Determination of the *m*-value for clamp association. The abscissa is the standard free energy change for the association of two clamps, determined as −*RTlnK*_*β*_2__, where T = 293 K and *K*_*β*_2__ is the equilibrium constant determined as discussed in the text.

### Sample preparation for FCS experiments

The assay buffer for all FCS experiments was 20 mM Tris-HCl (pH 7.5) containing variable concentrations of NaCl, KGlu, or glycine betaine (GB) as indicated in each experiment. For the experiments of Figs. 2A, 4A, 5A, and S1-S4, The stock labeled protein (concentration = 10-80 μM depending on prep) was first diluted to 200 nM using 20 mM Tris-HCl (pH 7.5), and this sample was further diluted to a final concentration of 1 nM and a final volume of 200 μL using 20 mM Tris-HCl (pH 7.5) and the desired concentration of salt or GB. Experiments as a function of protein concentration were carried out by mixing variable volumes of a 200 nM stock of labeled clamp, a 2 μM stock of labeled clamp, and a 2.5 M stock of KGlu or GB. All three solutions were prepared in 20 mM Tris-HCl (pH 7.5). Measurements were carried out immediately after mixing.

**Fig. 4:**
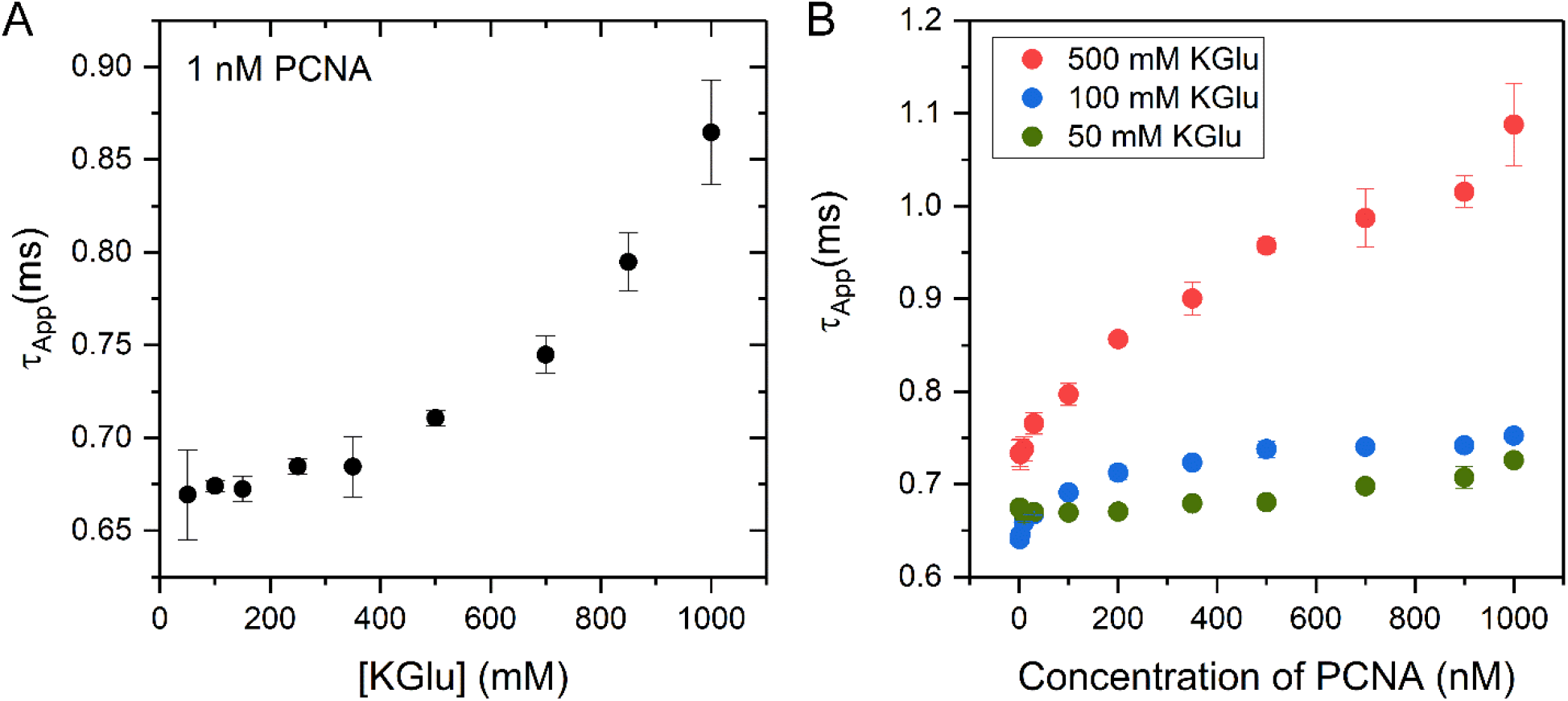
**(A)** Apparent diffusion times (τ_App_) obtained from the FCS decays measured with 1 nM solutions of PCNA-I181C-TMR in an assay buffer containing 20 mM Tris-HCl (pH 7.5) and a variable concentration of KGlu in the 50 mM – 1M range. **(B)** Apparent diffusion times (τ_App_) of solutions containing 1 nM PCNA-I181C-TMR and variable concentrations of unlabeled PCNA. The concentration specified in the abscissa represents the sum of the two. The assay buffer was 20 mM Tris-HCl (pH 7.5) containing variable concentrations of KGlu.

**Fig. 5:**
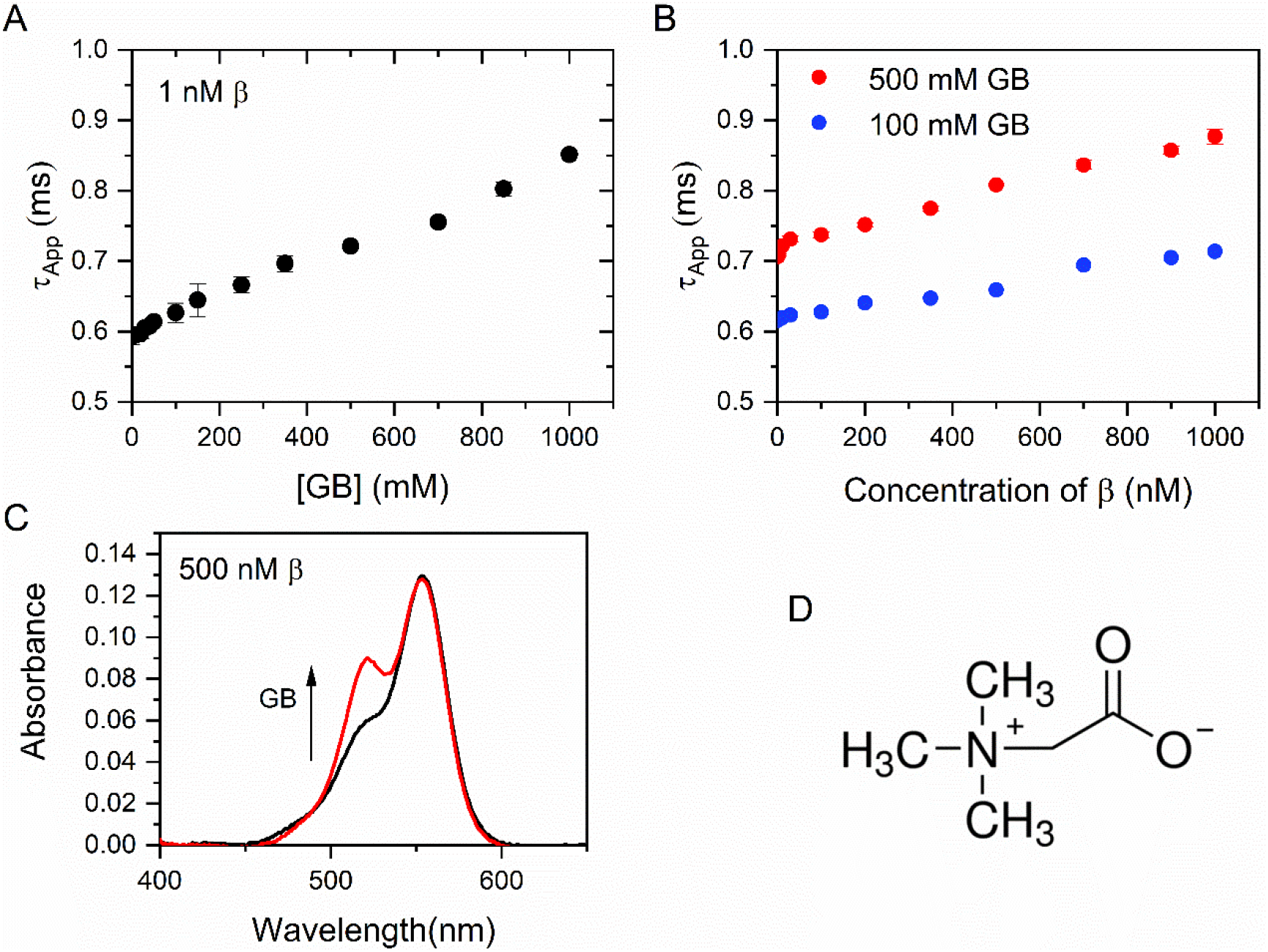
**(A)** Apparent diffusion times (τ_App_) obtained from the FCS decays measured with 1 nM solutions of β-I305C-TMR in an assay buffer containing 20 mM Tris-HCl (pH 7.5) and a variable concentration of GB as indicated in the abcissa. **(B)** Apparent diffusion times (τ_App_) of solutions containing 1 nM β-I305C-TMR and variable concentrations of unlabeled β. The concentration specified in the abscissa represents the sum of the two. The assay buffer was 20 mM Tris-HCl (pH 7.5) containing the indicated concentrations of GB. **(C)** Absorption spectra of 0.5 M β-I305C-TMR in Tris-HCl (no GB, in black) and in Tris-HCl containing 1 M GB (red) **(D)** Chemical structure of glycine betaine (GB).

### Analysis of FCS data and determination of τ_App_

The autocorrelation function for a monodisperse sample diffusing freely in solution is:

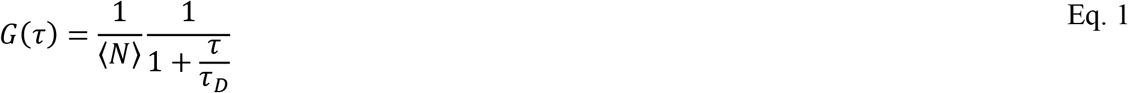

where τ is the correlation lag time, τ_D_ is the characteristic diffusion time, and 〈*N*〉^−1^ is the mean number of fluorescent particles in the observation volume. The characteristic diffusion time is related to the particle’s diffusion coefficient (*D*) as 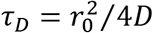, where r_0_ is the radial semi-axis of the Gaussian observation volume. The above equation is valid when the axial dimension of the observation volume is much larger than its radial dimension, which is the case in our instrument (10,38).

In the case of a polydisperse solution, the total autocorrelation function can be written as the sum of individual components weighted by the square of the brightness of each particle:

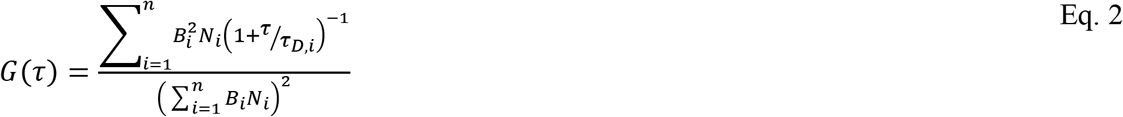

where, *n* represents the number of species present in solution, *N*_i_ represents the number of particles of species *i* present in the observation volume on average, *B*_i_ represents the brightness of species *i*, and τ_*D*,*i*_ represents its characteristic diffusion time (39,40).

The total autocorrelation decay of a mixture of oligomers (Eq. 2) is experimentally indistinguishable from the autocorrelation decay of a monodisperse sample (Eq. 1). Experimental data was therefore fitted with Eq. 1, which contains the least amount of fitting parameters (see previous publications from our group for an in-depth discussion) (9,10,38). Diffusion times obtained in this way are called apparent diffusion times (τ_App_), and depend on *N*_i_, *B*_i_ and τ_*D,i*_ (38).

### UV-VIS Absorption Measurements

UV-Vis absorption measurements were carried out in a Shimadzu UV-1700 PharmaSpec UV-Vis spectrophotometer. Solutions were contained in 1 cm-path length quartz microcuvettes (100 μL volume). Proteins were dissolved in 20 mM Tris-HCl buffer pH 7.2 with variable concentrations of KGlu or GB. The concentration of protein was kept constant at 1 μM for all UV-Vis experiments.

### Water-accessible surface area (ASA) calculations

ASA values were calculated from pdb files using the SurfRacer program (41) using a 1.4 Å radius for water and the Richards set of van der Waals radii. The crystallographic structure of β was obtained from the Protein Data Bank (pdb id: 1mmi). The pdb file for the dimer of rings was created in PyMol as detailed in the supplemental information file (Figure S5).

## RESULTS AND DISCUSSION

### KGlu prevents clamp dissociation and promotes clamp assembly

We previously reported the effect of NaCl on the dissociation equilibrium of β (10). Increasing NaCl concentration results in an increase in the equilibrium dissociation constant (*K*_d_) of the dimeric clamp and faster subunit dissociation kinetics. The dependence of *K*_d_ on NaCl concentration is consistent with simple charge-screening models; that is, the logarithm of *K*_d_ increases linearly with the square root of the molar concentration of salt (10). In addition, 1 nM β is stable as a dimer over 48 h in 50 mM NaCl buffer, but dissociates into monomers with a half-life of about 1 h in buffers containing 1 M NaCl (9). If salt effects on clamp dissociation equilibria are exclusively of electrostatic nature, any 1:1 electrolyte should affect the equilibrium constants in the same way. Here, we report on the oligomerization properties of β in KGlu, the primary cytoplasmic salt in *E. coli*. We employed the same strategy used previously to investigate clamp dissociation in NaCl, where fluorescence correlation spectroscopy experiments were carried out to determine the apparent diffusion times (τ_App_) of solutions of β in buffers of variable salt concentration (9,10). For a solution containing a single fluorescent species (i.e. one single diffusion coefficient), diffusion times are related to the diffusion coefficient of the fluorescent particle as

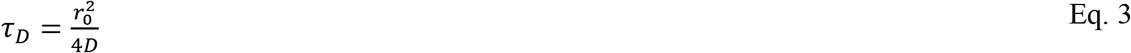

where *D* is the diffusion coefficient of the fluorescent particle and *r*_0_ is the radial semi-axis of the Gaussian observation volume (38,42). The experimental autocorrelation function of a solution containing a mixture of oligomeric species depends on the concentration, relative brightness, and diffusion coefficient of each particle in solution. Yet, although species of different size contribute to the total autocorrelation function, FCS decays usually fit well with a single diffusion time. This is because the diffusion coefficient of a spherical particle of radius *a* is proportional to *a*^−1^, and therefore to *V*^−1/3^, where *V* is the volume of the sphere. As a result, τ_D_ values are approximately proportional to the cube root of the number of protomers in the oligomeric protein, resulting in τ_D_ values that only increase by ~25% when the volume of the protein doubles. Experimentally, the autocorrelation decay of a mixture of dimers (D) and monomers (M) can be fitted with a single apparent diffusion time (*τ_M_* < *τ_App_* < *τ_D_*) that depends on the degree of dissociation of the dimeric protein. In general, the *τ_App_* of a mixture of labeled oligomers depends on the diffusion coefficients of the individual species, their fractional concentrations, and their relative brightness (38). We have previously reported the mathematical formalism that allows the calculation of the degree of dissociation of an oligomeric protein from measured τ_App_ values (38). The same formalism can be used to quantitate equilibrium dissociation and association constants between a broad variety of fluorescently labeled biopolymers, and will be used here to analyze glutamate-dependent clamp dissociation and association reactions.

Proteins for FCS were labeled with a fluorophore in each protein subunit as described in Materials and Methods and reported previously (9,10,43). Briefly, the surface-exposed amino acid Ile-305 of β was mutated to Cys, and the mutant protein was reacted with the maleimide derivative of tetramethylrhodamine (TMR). This procedure introduces a single TMR label in each subunit, resulting in a clamp with two fluorescent dyes placed at diametrically opposite positions (β-I305C-TMR, Fig. 1). These constructs are identical to the ones used in our previous work (10,33,34) and, importantly, the crystal structure of a closely related labeled clamp was previously solved and control experiments were carried out to rule out the potential adverse effects of mutations and fluorescent labeling on β structure and function (34).

FCS measurements were made on solutions of 1 nM β-I305C-TMR in an assay buffer containing 20 mM Tris-HCl (pH 7.5) and a variable concentration of NaCl or KGlu in the 50 mM – 1M range. Autocorrelation decays were acquired immediately after diluting a 200 nM stock solution of protein stored in 20 mM TRIS buffer without added salts into TRIS buffer containing variable concentrations of NaCl or KGlu. τ_App_ values were obtained from the analysis of the experimental autocorrelation decays as described in Materials and Methods and reported previously (10,44). Measured τ_App_ values decrease slightly at high NaCl concentrations, but much less than what is expected for complete dimer dissociation (i.e. a ~ 2^1/3^-fold reduction that would lead to τ_App_ ≈ 0.48 ms). This indicates a small degree of clamp dissociation during data acquisition (10 min) at high salt concentrations, consistent with our previous observations that 1 nM β is stable as a dimer over 48 h in 50 mM NaCl buffer, but dissociates into monomers with a half-life of about 1 h in buffers containing 1 M NaCl (9). Experiments with KGlu, however, gave markedly different results. Not only τ_App_ values do not decrease which increasing salt concentration (which would indicate clamp dissociation), but they increase steeply indicating a pronounced decrease in the diffusion coefficient of the protein particles. A control FCS experiment was performed with free TMR dye in 1 M KGlu-TRIS buffer to rule out the possibility that the observed changes in τ_App_ were due to an increase in the viscosity of the buffer. The diffusion coefficient of free TMR is independent of KGlu concentration in the 0-1M range within error, ruling out viscosity effects.

The increase in τ_App_ observed at high KGlu concentrations therefore indicates an increase in the radius of gyration of the protein, which suggests the formation of protein assemblies. The horizontal lines at the right side of Figs. 2A and 2B are placed at values of τ_App_ = τ_1_*n*^1/3^ and approximately indicate the diffusion times expected for protein assemblies containing *n* = 1, 2, … clamps. The diffusion time measured in 1 M KGlu-TRIS buffer indicates that 1 nM β assembles into structures containing an average of *c.a*. 3.5 clamps. Therefore, replacing NaCl by KGlu not only prevents clamp dissociation, but also promotes the association of two or more clamps into a protein assembly. τ_App_ values were measured immediately after diluting the protein stock into the buffer containing glutamate, and remained constant for over 1 hour indicating that equilibration is rapid and assemblies are kinetically stable. FCS experiments were then repeated replacing KGlu by NaGlu to dissect the contributions of the cation and the anion. The results of these experiments (see Fig. S1) are identical within experimental error, indicating that changes in τ_App_ are a consequence of replacing Cl^-^ by Glu^-^ in the buffer.

To rule out the possibility that the fluorescent tags induce or interfere with the formation of the assemblies, we carried out control experiments with clamps labeled with Alexa 546 instead of TMR (β-I305C-A546), and with clamps labeled with TMR but in different positions within the clamp. Alexa 546 is significantly bulkier than TMR (see Fig. S2) and it has a reduced tendency to aggregate in aqueous solution compared to TMR due to the sulfonic acid moieties present in the Alexa dyes and absent in TMR (43). FCS experiments with β-I305C-A546 in buffers containing KGlu show the same τ_App_ trends observed with the TMR-labeled clamp (Fig. S2), indicating that the dyes do not interfere with or induce the assembly process. Moreover, the same results were obtained in experiments using clamps labeled in different positions (Fig. S3). These control experiments give us confidence that the KGlu-dependent changes in diffusion are indeed due to the inherent tendency of the clamp to assemble in these conditions. To our knowledge, association of sliding clamps was only observed previously in ion mobility mass spectrometry experiments, where signals corresponding to two β clamps (i.e. a dimer of dimers) were observed (45).

### Clamp assembly is governed by the law of mass action

Results so far indicate the formation of protein assemblies when KGlu is added to a solution of β. To evaluate whether protein assembly is reversible, we repeated FCS experiments in the reverse direction, that is, from high to low glutamate concentration. A 200 nM stock solution of β in 1M KGlu buffer was prepared, and 200-fold dilutions were performed using TRIS buffer containing variable concentrations of glutamate in the 10 nM-1M range. Results (Fig. S4) indicate that protein assemblies dissociate when the concentration of glutamate in the buffer decreases, and apparent diffusion times depend only on KGlu concentration, and not on whether proteins were initially stored in high or no KGlu. Therefore, protein assembly is a reversible process, and that the size of the assemblies is solely determined by the final Glu^-^ concentration in the buffer.

The results of Fig. 2A are consistent with the formation of protein assemblies with an equilibrium association constant that increases with increasing KGlu concentration. The formation of assemblies at protein concentrations as low as 1 nM suggests that association constants are of the order of 1 nM^−1^ or greater at high KGlu concentration. Therefore, at a given KGlu concentration, increasing the concentration of protein should result in a distribution of assemblies with a greater average size. To test this prediction, FCS experiments were repeated using solutions of β in the 1nM – 1 μM range. The amplitude of the FCS decay is inversely proportional to the concentration of fluorescently labeled protein, which restricts FCS measurements to the ca. 1 nM-100 nM range. To determine τ_App_ values at higher protein concentrations, the concentration of labeled protein was kept constant at 1 nM while the total protein concentration (which determines its equilibrium properties) was adjusted by adding unlabeled clamps. This design also reduces the average number of fluorescent labels per protein assembly, which avoids the undesirable formation of very bright assemblies that would mask emission from singly labeled monomers. Due to the dimeric nature of β, each clamp carries two fluorescent dyes placed at diametrically opposite positions (Fig. 1). We have previously determined that the exchange of labeled and unlabeled subunits within a clamp is slow and requires incubation over 24 h (9). Here, FCS decays were acquired immediately after mixing the labeled and unlabeled proteins, so clamps are not expected to exchange their subunits in the timescale of these experiments. Measured τ_App_ values do not depend on protein concentration at low concentration of glutamate (10 mM KGlu, Fig. 2B), and are equal to the values measured in 50 mM NaCl. This indicates that clamps are stable as dimeric rings, and 10 mM KGlu is not enough to promote clamp association in the 1 nM-1 μM protein concentration range. Clamp association becomes significant at KGlu concentrations of the order of hundreds of millimolar, and the average size of the protein assemblies that exist in equilibrium at each KGlu concentration increases with increasing protein concentration, as expected from the law of mass action. The horizontal bars at the right side of Fig. 2B are placed at a value of τ_App_ = 0.6*n*^1/3^, where 0.6 ms is the diffusion time of the β ring (*n* = 1). These τ_App_ values are the expected diffusion times for an assembly of *n* rings provided that all diffusion particles can be regarded as spherical and that the volume of the assembly scales linearly with the number of rings. In addition, TMR self-quenching at high protein and high glutamate concentrations (see below) adds uncertainty to these calculated *n* values, which should be only taken as rough estimates.

### Clamp assemblies involve face-to-face interactions between adjacent rings

Fig. 2C shows the absorption visible spectrum of 1 μM β-I305C-TMR in TRIS buffer containing 50 mM – 1 M KGlu. The absorption spectrum of the protein solution is consistent with the absorption spectrum of the free dye at low KGlu concentrations, but changes dramatically at KGlu concentrations higher than ca. 250 mM. The increase in absorbance at the blue shoulder of the spectrum observed at higher concentrations of KGlu is a well-known feature of H-type rhodamine dimers, which are characterized by a plane-to-plane stacking geometry of two dye molecules separated by van der Waals distances (43,46). These results indicate that two molecules of rhodamine are brought together in close proximity at glutamate concentrations where measured τ_App_ values indicate the formation of protein assemblies involving several rings (see τ_App_ values of 1 μM β at different KGlu concentrations, Fig. 2B). The geometric requirements for observing these distinct spectroscopic features are quite stringent (47,48), so the strong absorption signal measured at 520 nm indicates efficient formation of rhodamine dimers, which can only be explained if two clamps assemble in a face-to-face configuration. Dimerization of rhodamine molecules in TRIS buffer occurs only at concentrations greater than 100 mM (43,46), and 1 μM free TMR in 1 M KGlu TRIS buffer has the spectrum of the monomeric dye (not shown). Therefore, the formation of H-type rhodamine dimers in a 1 μM solution of β-I305C-TMR can only be explained by the association of two clamps that bring the two dyes together at the required distance and orientation. A model of a possible clamp-clamp interaction that is consistent with rhodamine dimerization is shown in Fig. 2D. This model was created using the crystallographic structure of β (pdb id: 1mmi) and coordinates for tetramethylrhodamine-5-maleimide extracted from the x-ray crystal structure of tetramethylrhodamine-5-maleimide-actin (PDBeChem id: RHO). PyMOL was used to replace Ile-305 by cys, and to create a bond between cys and tetramethylrhodamine-5-maleimide. As shown in Fig. 2D, face-to-face interactions between two clamps would allow the rhodamine dyes to stack closely in the conformation required to explain the strong 520 nm absorption band. We note that the formation of a second dimer of rhodamine is also possible on the opposite side of the clamp (not shown for clarity). PyMOL models suggest that this is the only orientation of the protein rings that allow the rhodamine dyes to interact in this manner. However, diffusion times at high glutamate concentrations are consistent with protein assemblies larger than a dimer, indicating that other modes of interaction that may not result in the formation of rhodamine dimers are also possible (see below).

### Molecular mechanism of glutamate-induced clamp association

Many salts (49,50) and nonelectrolytes such as sugars (51), polyols (52,53), and amino acids (54), are known to stabilize globular proteins and to reduce their solubility. *In vitro*, replacing Cl^-^ by Glu^-^ stabilizes folded proteins and stabilizes protein-nucleic acid complexes (18–26). These effects are only observed at high (> 100 mM) concentrations of these added agents (henceforth called solutes), indicating that their interactions with the surface of proteins are weak and non-specific (55,56). Excluded volume effects due to molecular crowding have been proposed as mechanisms for osmolyte-induced protein stability (57–59), but contributions are significant only for larger solutes such as high molecular weight PEGs (20,60,61). Alternatively, the effect of osmolites on protein stabilization and solubility can be rationalized in terms of the relative binding affinities of the protein surface for water or for the solute. Solutes that are able to displace water from the protein surface due to their greater affinity for the exposed protein functional groups are known to destabilize the native state of proteins. Likewise, a greater binding affinity of the protein surface for water than for the solute results in a net exclusion of the solute from the protein surface, which has a stabilizing effect (55,56,62).

In the context of this interpretation, the effect of solutes on the chemical potential of the protein can be described using three-component thermodynamic theory where water, protein and solute are taken into account implicitly (56). Whether addition of a solute will promote a macromolecular process such as protein unfolding or protein assembly depends on the effect of the solute on the standard free energy change of the process (ΔG°). Let us first consider the association of two β clamps to form a dimer: 2*β* ⇌ *β*_2_, with an equilibrium association constant *K*_*β*_2__ (Fig. 3A). This association is not thermodynamically favorable in water (i.e. no KGlu added), so 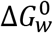, the change in standard free energy for the formation of β_2_ in water, is positive (Fig. 3A). To rationalize the sign of ΔG° in a solution containing glutamate at a given concentration 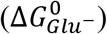, a thermodynamic cycle can be constructed in which two *β* clamps or the dimer *β*_2_ are hypothetically transferred from water to the glutamate solution (vertical lines in Fig. 3A). The standard free energy changes for these processes (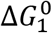 and 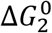) measure the extent by which the chemical potential of the protein increases or decreases in a hypothetical process in which the protein is transferred from water to the solution containing the solute. From Fig. 3A, 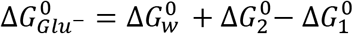, and therefore the formation of *β*_2_ will be favorable in glutamate buffer (i.e. 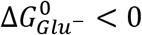) if 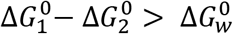.

The accessible surface area of a protein can be thought of as being composed of sites that bind water molecules. When the protein is transferred from water to a solution containing a given concentration of solute, the change in free energy is determined by the relative affinity of these sites for water and for the solute. Favorable interactions between the solute and the protein surface result in a reduction of the chemical potential of the protein (negative values of 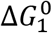 and 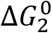). Similarly, if interactions are stronger for water than for the solute, the solute will be excluded from the protein surface and both 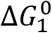 and 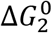 will be positive. For a given solute at a given concentration, the extent by which addition of a solute to water increases or decreases the chemical potential of the protein (measured by the vertical lines in Fig. 3A) depends on the protein’s accessible surface area. Cheng et al quantified the interactions between KGlu and model compounds containing six protein functional groups: sp^3^ carbon, sp^2^ carbon, amide oxygen, carboxylate oxygen, amide nitrogen, and cationic nitrogen (20). Interactions between KGlu and the first four functional groups are unfavorable (compared to water as a protein ligand), while interactions with amide and cationic nitrogen are favorable. The surface of a native protein is about 50% aliphatic or aromatic hydrocarbon, and about 20% amide oxygen and nitrogen (63). Most amide groups on the surface of proteins are in the peptide backbones, with a O:N available surface area ratio of about 2.4:1 (63). In addition, the absolute value of the interaction potential of KGlu with amide oxygen (which is unfavorable) is greater than the corresponding value for the favorable interaction of KGlu with amide nitrogen. Therefore, the net interaction between KGlu and the amide group is unfavorable. Since carbon and amide atoms make up a significant fraction of the protein surface, KGlu is expected to be highly excluded, and in terms of the quantities defined in Fig. 3A, 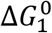 and 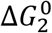 are both predicted to be positive. The formation of β_2_ is expected to occlude a fraction of the accessible surface area of the ring, so the unfavorable interactions between KGlu and the protein surface are predicted to be more significant for the two isolated rings than for the dimer. These arguments lead to the conclusion that 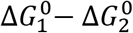 is always positive for processes that occlude surface, and if the difference is significant (i.e. greater than 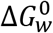), clamp association can become thermodynamically favored in the presence of glutamate (i.e. 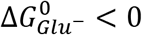).

We therefore conclude that unfavorable interactions of KGlu or NaGlu with the functional groups on the protein surface satisfactorily explain the marked effects of glutamate concentration on the equilibrium constant for the formation of β_2_ (*K*_*β*_2__). If this interpretation is correct, glutamate effects should be observed in experiments with other clamps and in general, with other proteins that can bury a significant fraction of their surface by associating. In addition, this model predicts that other solutes that are known to be excluded from the protein surface should promote clamp association as well. Finally, the effect of KGlu on *K*_*β*_2__ should be quantitatively consistent with the amount of accessible surface area excluded by the interaction between the clamps and the interaction potentials between KGlu and the protein functional groups. These three points are addressed individually in the sections below.

### KGlu effects are not specific to β

To assess whether glutamate effects are due to specific interactions between this solute and β, FCS experiments were repeated replacing β by the proliferating cell nuclear antigen (PCNA) sliding clamp from *S. cerevisiae* (See Fig. 1). The three-dimensional structures of PCNA and β are very similar despite the facts that PCNA is a homotrimer, and the two proteins show no sequence homology (64). Proteins for FCS were labeled with a TMR fluorophore in each protein subunit as described in Materials and Methods and reported previously (9). FCS experiments were performed with 1 nM solutions of labeled PCNA (PCNA-I181C-TMR, Fig. 1) in buffers containing variable concentrations of KGlu, and apparent diffusion times were calculated from the experimental autocorrelation decays as described above for β. As in the case of β, FCS results are consistent with the formation of PCNA assemblies with an association equilibrium constant that increases with KGlu concentration (Fig. 4). However, we did not observe changes in the UV-VIS absorption spectrum of PCNA-I181C-TMR in conditions where FCS experiments suggest the formation of clamp assemblies involving several rings (1 μM PCNA, KGlu > 500 mM). This is not entirely surprising given how stringent the geometrical requirements for the formation of H-dimers are, and suggests that clamp-clamp interactions do not result in the placement of TMR dyes at the distance and geometry required for efficient coupling of the rhodamine rings.

### KGlu effects on association equilibria are quantitatively consistent with KGlu-protein interactions

If the experimental results of Figs. 2A and 2B are indeed due to unfavorable interactions between KGlu and the functional groups on the protein surface, the effect of KGlu concentration on the association equilibrium constant between two clamps (*K*_*β*_2__) should be quantitatively consistent with the change in accessible surface area (ASA) that results from the formation of β_2_. The interaction potentials between KGlu and six different protein functional groups have been measured experimentally (20), and these values can be used to make quantitative predictions of the effect of KGlu concentration on the equilibrium constants of processes that change the protein’s water-accessible surface area. For example, Cheng et. al. studied the effect of KGlu on the unfolding of the native state of the protein NTL9, and demonstrated that these interaction potentials result in predicted *m*-values (defined as the derivative of ΔG of unfolding with respect to solute concentration (65)) that are remarkably similar to the experimentally determined values (19,20).

In analogy to the *m*-value defined for protein unfolding, we considered the change in −RT*lnK*_*β*_2__ with respect to KGlu concentration. Note that concentrations are now given in molal units (m) as required in the analysis using interaction potentials (see below). Values of *K*_*β*_2__ were determined from the τ_app_ values measured in conditions where the concentrations of trimers, tetramers, and higher order oligomers are negligible ([KGlu] ≈ 0-500 mM). Details are given in the supplemental information file (see “Calculation of *K*_*β*_2__”), and results are shown in Fig. 3B. The slope of the straight line of Fig. 3B (−5.1 ± 0.7 kcal mol^−1^ m^−1^) is the experimental *m*-value for the equilibrium 2*β* ⇌ *β*_2_ at 20°C. We next calculate the *m*-value predicted for clamp association under the assumption that changes in *K*_*β*_2__ are due to chemical interactions between KGlu and functional groups at the protein surface. A quantitative agreement between the experimental and theoretical *m*-values would provide strong support for the model used to interpret glutamate-dependent variations in *K*_*β*_2__. Following the formalism used by Cheng. et. al. to analyze the effect of KGlu on protein unfolding equilibria, the theoretical *m*-value was calculated as (20):

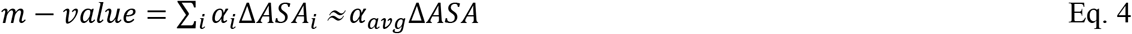

where, α_i_ represents the interaction potential per unit of ASA between KGlu and each of the protein’s functional groups, and ΔASA_i_ is the change in accessible surface area of each individual functional group during the process under consideration (i.e. the formation of β_2_) (20). The exact structure of β_2_ is unknown, so a calculation atom by atom is not possible. Instead, we will assess whether the change in total accessible surface area required to justify the experimental *m*-value is consistent with a reasonable structural model of β_2_. Assuming a constant average value of α_avg_ = 1.2 cal mol^−1^ m^−1^ Å^−2^, the experimental *m*-value requires a change in the accessible surface area of ΔASA = −4,250 ± 600 Å^2^ (Eq. 4). The value of α_avg_ was estimated from the values of α_i_ published by Cheng et al. (20) and the fact that approximately half of the surface of native proteins is aliphatic carbon (α_i_ = 1.34 cal mol^−1^ m^−1^ Å^−2^) and about 20% is amide oxygen (α_i_ = 0.76 cal mol^−1^ m^−1^ Å^−2^) and nitrogen (α_i_ = −0.39 cal mol^−1^ m^−1^ Å^−2^) with an O:N ASA ratio of approximately 2.4:12 (20,63). To evaluate whether a ΔASA of −4,250 Å^2^ is reasonable, we created model structures of β_2_ using PyMOL and calculated their ASAs computationally. Model dimers were created from two initially overlapping copies of β (pdb entry: 1mmi) by displacing one of the clamps a given distance (*d*) along the axis normal to the plane of the ring and then rotating one of the clamps 180° to create a structure compatible with rhodamine dimerization (see Supplemental Information for details). The pdb files of the model dimers were then analyzed using the Surface Racer program as described in Materials and Methods, and ΔASA values for the formation of β_2_ were then calculated as *ASA*_*β*_2__ – 2*ASA_β_*. Results were −3230 Å^−2^, −4052 Å^−2^, and −4850 Å^−2^ for displacements of *d*=34 Å, *d*=33 Å and *d*=32 Å, respectively (see Fig. S5). Therefore, we conclude that dimerization of β results in a reduction in accessible surface area that is quantitatively consistent with the observed glutamate-dependent variations in *K*_*β*_2__. In other words, our data can be quantitatively explained in terms of chemical unfavorable interactions between KGlu and the functional groups of the surface of the clamp.

### Clamp association is not specific to Glu^-^

If glutamate effects are indeed due to unfavorable interactions between glutamate and the functional groups at the protein surface, other chemical compounds that are also preferentially excluded from the surface of proteins should promote clamp assembly as well. To test this prediction, we carried out FCS experiments replacing KGlu by glycine betaine (GB, Fig. 5D), a zwitterionic compound that accumulates intracellularly when *E.coli* and most other living cells are exposed to high concentrations of extracellular solutes (30–32). At the molecular level, the effectiveness of GB as an osmolyte has been interpreted as arising from its unfavorable interactions (compared with water) with anionic and amide oxygen atoms at the protein surface (66). As in the case of KGlu, the interaction potentials per unit of ASA (α_i_) for GB-protein interactions are positive for aliphatic carbon and for carboxylate and amide oxygen, and negative for cationic and amide nitrogen (66). Given the composition of the protein surface, it is expected that interactions between GB and the protein functional groups will be mostly unfavorable, resulting in a preferential exclusion of GB from the surface.

Fig. 5A shows the diffusion times extracted from the autocorrelation decays acquired in FCS experiments made on solutions of 1 nM β-I305C-TMR in an assay buffer containing 20 mM Tris-HCl (pH 7.5) and a variable concentration of GB in the 50 mM – 1M range. As in the case of KGlu, τ_App_ values are consistent with the formation of protein assemblies with an average size that increases with protein concentration (Fig. 5B). The absorption spectra of solutions of 0.5 μM β-I305C-TMR in GB-TRIS buffers (Fig. 5C) indicate the formation of rhodamine dimers at high GB concentrations, suggesting the same type of clamp-clamp interactions observed with KGlu. While both GB and Glu promote clamp association, their quantitative effect on the corresponding association constants is somewhat different. This is not surprising because although both compounds are predicted to be preferentially excluded from the protein surface, their α_i_ values differ. For GB, interactions with aliphatic carbon, which represents about half of the protein surface, are almost neutral (small positive α_i_), while interactions with amide and carboxylate oxygen are highly unfavorable (large positive α_i_). In contrast, KGlu is predicted to interact very unfavorably with aliphatic and aromatic carbon, and to a lesser extent, with amide oxygen.

### Possible modes of assembly

The characteristic spectrum of rhodamine dimers observed in the visible absorption spectrum of 1 μM β-I305C-TMR in high KGlu (≥ 350 mM) can only be explained if two clamps interact in the configuration shown in Fig. 2D. In these protein constructs, the dye is covalently bound to a cysteine residue that replaces I305 (Fig. 1), which is located on the face of the clamp opposite to the one involved in the interaction between β and the γ clamp loader complex (67). This, however, cannot be the only mode of clamp-clamp interaction because it would result in the formation of dimers of rings, and FCS results (Fig. 2B) are consistent with a continuous assembly of rings into larger structures. While the spectroscopic signature of Fig. 2C is unambiguous, FCS experiments (Fig. 2B) suggest that 1 μM β associates as dimers of rings at KGlu concentrations lower than the 350 mM needed to observe strong coupling between two rhodamines. This suggests that the initial interaction may be different from the one shown in Fig. 2D. In addition, FCS experiments with 1 nM β show significant changes in τ_App_ without changes in the total fluorescence intensity. Rhodamine dimers are not fluorescent, so their formation is expected to result in significant quenching and a concomitant reduction in measured intensity. The fact that 1 nM β associates tightly in high glutamate without fluorescence quenching favors a model in which the most thermodynamically favorable interactions do not bring the two dyes in close proximity. Evidence for different modes of interaction with significantly different equilibrium constants also comes from Fig. 2B. While τ_App_ values increase steeply initially consistent with association constants of the order of 1 nM^−1^ and higher, further changes are not as pronounced and indicate weaker clamp-clamp interactions. In summary, our data is consistent with clamp-clamp interactions that bury *c.a*. 4000 Å^−2^, a value that is consistent with the face-to-face interaction of two rings separated by about 33 Å. Although the spectroscopic signature observed in the visible spectrum of β-I305C-TMR provide strong support for interactions between the faces proximal to I305 (Fig. 2D), results overall point to a more complex scenario involving other modes of interactions with varied affinities. The large surface buried in face-to-face interactions results in a strong dependence of the affinity constants with glutamate and glycine betaine concentrations.

### Biological Implications

The number of β-clamps has been estimated to be about 300 - 350 molecules per cell (68,69). Using an estimate of 1 fL (10^−15^ L) for an average cell volume, the concentration of β-clamps is on the order of 500 - 600 nM in *E. coli* (70,71). At a normal cellular glutamate concentration of 100 mM, data in Figure 2B indicate that clamps would exist as a mixture of monomers and dimers with dimers predominating inside cells. Stressors that increase the intracellular glutamate concentration would increase the population of dimers and lead to formation of some trimeric species. What are the implications for clamp oligomerization inside the cell? Oligomerization could sequester clamps and reduce the effective intracellular concentration of clamps available for DNA metabolism. The symmetry of the clamp is such that it has two distinct faces like the head and tail sides of a coin. Proteins that bind the clamp, including the clamp loader and DNA polymerase, bind the same site on one of the faces. If clamps were to dimerize in a ‘head-to-tail’ fashion as suggested by the analysis above, then the binding face of one clamp would be sequestered and the effective concentration of available clamps would be reduced by a factor of two. These dimers could be extended further by adding additional clamps in a head-to-tail fashion. Sequestering binding faces could potentially provide a mechanism for slowing DNA metabolism under times of stress. On the other hand, if clamps were to dimerize in a ‘tail-to-tail’ fashion as in Figure 2B, then the ‘heads’ side to which proteins bind would still be available for binding, and the effective concentration of available clamps would remain the same. This clamp stacking argument assumes that oligomerization does not affect clamp activity in some way other than physically blocking binding interactions. But, it is also possible that oligomerization in either the head-to-tail or tail-to-tail orientation alters clamps activity in some way to interfere with productive interactions with clamp binding partners. This would effectively sequester clamps to reduce the number of clamps available for DNA metabolism.

Clamp oligomerization could have a completely different function and potentially serve to stabilize the ring structure of individual clamps. Ring-shaped clamps are loaded onto DNA by clamp loaders. Dissociation of the rings into monomers would prevent them from being loaded onto DNA by the bacterial clamp loader. Competing factors could exist inside cells that have differential effects on the stability of the ring structure. If conditions exist that promote the dissociation of rings into individual monomers, oligomerization of the clamps could counteract these conditions and stabilize the rings to maintain their availability for DNA metabolism.

## REFERENCES

1. Hedglin, M., Kumar, R. and Benkovic, S.J. (2013) Replication clamps and clamp loaders. Cold Spring Harb Perspect Biol, 5, a010165.

2. Johnson, A. and O’Donnell, M. (2005) Cellular DNA replicases: Components and dynamics at the replication fork. Annu Rev Biochem, 74, 283–315.

3. Bloom, L.B. (2006) Dynamics of loading the Escherichia coli DNA polymerase processivity clamp. Crit Rev Biochem Mol Biol, 41, 179–208.

4. Jeruzalmi, D., O’Donnell, M. and Kuriyan, J. (2002) Clamp loaders and sliding clamps. Curr Opin Struct Biol, 12, 217–224.

5. Krishna, T.S., Kong, X.P., Gary, S., Burgers, P.M. and Kuriyan, J. (1994) Crystal structure of the eukaryotic DNA polymerase processivity factor PCNA. Cell, 79, 1233–1243.

6. Moarefi, I., Jeruzalmi, D., Turner, J., O’Donnell, M. and Kuriyan, J. (2000) Crystal structure of the DNA polymerase processivity factor of T4 bacteriophage. J Mol Biol, 296, 1215–1223.

7. Kong, X.P., Onrust, R., O’Donnell, M. and Kuriyan, J. (1992) 3-Dimensional Structure of the β-Subunit of E. coli DNA Polymerase-III Holoenzyme - a Sliding DNA Clamp. Cell, 69, 425–437.

8. Yao, N., Turner, J., Kelman, Z., Stukenberg, P.T., Dean, F., Shechter, D., Pan, Z.Q., Hurwitz, J. and O’Donnell, M. (1996) Clamp loading, unloading and intrinsic stability of the PCNA, beta and gp45 sliding clamps of human, E. coli and T4 replicases. Genes Cells, 1, 101–113.

9. Binder, J.K., Douma, L.G., Ranjit, S., Kanno, D.M., Chakraborty, M., Bloom, L.B. and Levitus, M. (2014) Intrinsic stability and oligomerization dynamics of DNA processivity clamps. Nuc Acids Res, 42, 6476.

10. Purohit, A., England, J.K., Douma, L.G., Tondnevis, F., Bloom, L.B. and Levitus, M. (2017) Electrostatic Interactions at the Dimer Interface Stabilize the E-coli beta Sliding Clamp. Biophys J, 113, 794–804.

11. Lo Nostro, P. and Ninham, B.W. (2012) Hofmeister phenomena: an update on ion specificity in biology. Chem Rev, 112, 2286–2322.

12. Collins, K.D. (2004) Ions from the Hofmeister series and osmolytes: effects on proteins in solution and in the crystallization process. Methods, 34, 300–311.

13. Zhang, Y. and Cremer, P.S. (2006) Interactions between macromolecules and ions: The Hofmeister series. Curr Opin Chem Biol, 10, 658–663.

14. Bennett, B.D., Kimball, E.H., Gao, M., Osterhout, R., Van Dien, S.J. and Rabinowitz, J.D. (2009) Absolute metabolite concentrations and implied enzyme active site occupancy in Escherichia coli. Nat Chem Biol, 5, 593–599.

15. Richey, B., Cayley, D.S., Mossing, M.C., Kolka, C., Anderson, C.F., Farrar, T.C. and Record, M.T., Jr. (1987) Variability of the intracellular ionic environment of Escherichia coli. Differences between in vitro and in vivo effects of ion concentrations on protein-DNA interactions and gene expression. J Biol Chem, 262, 7157–7164.

16. Cayley, S., Lewis, B.A., Guttman, H.J. and Record, M.T., Jr. (1991) Characterization of the cytoplasm of Escherichia coli K-12 as a function of external osmolarity. Implications for protein-DNA interactions in vivo. J Mol Biol, 222, 281–300.

17. McLaggan, D., Naprstek, J., Buurman, E.T. and Epstein, W. (1994) Interdependence of K+ and glutamate accumulation during osmotic adaptation of Escherichia coli. J Biol Chem, 269, 1911–1917.

18. Arakawa, T. and Timasheff, S.N. (1984) The mechanism of action of Na glutamate, lysine HCl, and piperazine-N,N’-bis(2-ethanesulfonic acid) in the stabilization of tubulin and microtubule formation. J Biol Chem, 259, 4979–4986.

19. Sengupta, R., Pantel, A., Cheng, X., Shkel, I., Peran, I., Stenzoski, N., Raleigh, D.P. and Record, M.T., Jr. (2016) Positioning the Intracellular Salt Potassium Glutamate in the Hofmeister Series by Chemical Unfolding Studies of NTL9. Biochemistry, 55, 2251–2259.

20. Cheng, X., Guinn, E.J., Buechel, E., Wong, R., Sengupta, R., Shkel, I.A. and Record, M.T., Jr. (2016) Basis of Protein Stabilization by K Glutamate: Unfavorable Interactions with Carbon, Oxygen Groups. Biophys J, 111, 1854–1865.

21. Leirmo, S., Harrison, C., Cayley, D.S., Burgess, R.R. and Record, M.T., Jr. (1987) Replacement of potassium chloride by potassium glutamate dramatically enhances protein-DNA interactions in vitro. Biochemistry, 26, 2095–2101.

22. Deredge, D.J., Baker, J.T., Datta, K. and Licata, V.J. (2010) The glutamate effect on DNA binding by pol I DNA polymerases: osmotic stress and the effective reversal of salt linkage. J Mol Biol, 401, 223–238.

23. Kozlov, A.G., Shinn, M.K., Weiland, E.A. and Lohman, T.M. (2017) Glutamate promotes SSB protein-protein Interactions via intrinsically disordered regions. J Mol Biol, 429, 2790–2801.

24. Menetski, J.P., Varghese, A. and Kowalczykowski, S.C. (1992) The physical and enzymatic properties of Escherichia coli recA protein display anion-specific inhibition. J Biol Chem, 267, 10400–10404.

25. Overman, L.B., Bujalowski, W. and Lohman, T.M. (1988) Equilibrium binding of Escherichia coli single-strand binding protein to single-stranded nucleic acids in the (SSB)65 binding mode. Cation and anion effects and polynucleotide specificity. Biochemistry, 27, 456–471.

26. Overman, L.B. and Lohman, T.M. (1994) Linkage of pH, anion and cation effects in protein-nucleic acid equilibria. Escherichia coli SSB protein-single stranded nucleic acid interactions. J Mol Biol, 236, 165–178.

27. Record, M.T., Jr., Courtenay, E.S., Cayley, S. and Guttman, H.J. (1998) Biophysical compensation mechanisms buffering E. coli protein-nucleic acid interactions against changing environments. Trends Biochem Sci, 23, 190–194.

28. Burg, M.B. and Ferraris, J.D. (2008) Intracellular organic osmolytes: function and regulation. J Biol Chem, 283, 7309–7313.

29. Yancey, P.H., Clark, M.E., Hand, S.C., Bowlus, R.D. and Somero, G.N. (1982) Living with water stress: evolution of osmolyte systems. Science, 217, 1214–1222.

30. Cayley, S. and Record, M.T. (2003) Roles of cytoplasmic osmolytes, water, and crowding in the response of Escherichia coli to osmotic stress: Biophysical basis of osmoprotection by glycine betaine. Biochemistry, 42, 12596–12609.

31. Kempf, B. and Bremer, E. (1998) Uptake and synthesis of compatible solutes as microbial stress responses to high-osmolality environments. Arch Microbiol, 170, 319–330.

32. Sakamoto, A. and Murata, N. (2002) The role of glycine betaine in the protection of plants from stress: clues from transgenic plants. Plant Cell Environ, 25, 163–171.

33. Douma, L.G., Yu, K.K., England, J.K., Levitus, M. and Bloom, L.B. (2017) Mechanism of opening a sliding clamp. Nuc Acids Res, 45, 10178–10189.

34. Paschall, C.O., Thompson, J.A., Marzahn, M.R., Chiraniya, A., Hayner, J.N., O’Donnell, M., Robbins, A.H., McKenna, R. and Bloom, L.B. (2011) The Escherichia coli Clamp Loader Can Actively Pry Open the beta-Sliding Clamp. J Biol Chem, 286, 42704–42714.

35. Thompson, J.A., Marzahn, M.R., O’Donnell, M. and Bloom, L.B. (2012) Replication Factor C Is a More Effective Proliferating Cell Nuclear Antigen (PCNA) Opener than the Checkpoint Clamp Loader, Rad24-RFC. J Biol Chem, 287, 2203–2209.

36. Anderson, S.G., Thompson, J.A., Paschall, C.O., O’Donnell, M. and Bloom, L.B. (2009) Temporal correlation of DNA binding, ATP hydrolysis, and clamp release in the clamp loading reaction catalyzed by the Escherichia coli gamma complex. Biochemistry, 48, 8516–8527.

37. Ranjit, S. and Levitus, M. (2012) Probing the Interaction Between Fluorophores and DNA Nucleotides by Fluorescence Correlation Spectroscopy and Fluorescence Quenching. Photochem Photobiol, 88, 782–791.

38. Kanno, D.M. and Levitus, M. (2014) Protein Oligomerization Equilibria and Kinetics Investigated by Fluorescence Correlation Spectroscopy: A Mathematical Treatment. J Phys Chem B, 118, 12404–12415.

39. Thompson, N.L. (1991) In Lakowicz, J. R. (ed.), Topics in Fluorescence Spectroscopy. Plenum Press, New York, Vol. 1, pp. 337–355.

40. Muller, J.D., Chen, Y. and Gratton, E. (2003) Fluorescence correlation spectroscopy. Method Enzymol, 361, 69–92.

41. Tsodikov, O.V., Record, M.T., Jr. and Sergeev, Y.V. (2002) Novel computer program for fast exact calculation of accessible and molecular surface areas and average surface curvature. J Comput Chem, 23, 600–609.

42. Elson, E.L. (2011) Fluorescence correlation spectroscopy: past, present, future. Biophys J, 101, 2855–2870.

43. Donaphon, B., Bloom, L.B. and Levitus, M. (2018) Photophysical characterization of interchromophoric interactions between rhodamine dyes conjugated to proteins. Methods Appl Fluoresc, 6, 045004.

44. Serban, A.J., Breen, I.L., Bui, H.Q., Levitus, M. and Wachter, R.M. (2018) Assembly-disassembly is coupled to the ATPase cycle of tobacco Rubisco activase. J Biol Chem, 293, 19451–19465.

45. Koleva, B.N., Gokcan, H., Rizzo, A.A., Lim, S., Fouque, K.J.D., Choy, A., Liriano, M.L., Fernandez-Lima, F., Korzhnev, D.M., Cisneros, G.A. et al. (2019) Dynamics of the E. coli beta-Clamp Dimer Interface and Its Influence on DNA Loading. Biophys J, 117, 587–601.

46. Arbeloa, I.L. and Ojeda, P.R. (1982) Dimeric States of Rhodamine-B. Chem Phys Lett, 87, 556–560.

47. Kemnitz, K. and Yoshihara, K. (1991) Entropy-Driven Dimerization of Xanthene Dyes in Nonpolar Solution and Temperature-Dependent Fluorescence Decay of Dimers. J Phys Chem, 95, 6095–6104.

48. Kasha, M., Rawls, H.R. and El-Bayoumi, M.A. (1965) The Exciton Model in Molecular Spectroscopy. Pure Applied Chemistry, 11, 371–392.

49. Arakawa, T., Bhat, R. and Timasheff, S.N. (1990) Why preferential hydration does not always stabilize the native structure of globular proteins. Biochemistry, 29, 1924–1931.

50. Dempsey, C.E., Mason, P.E., Brady, J.W. and Neilson, G.W. (2007) The reversal by sulfate of the denaturant activity of guanidinium. J Am Chem Soc, 129, 15895–15902.

51. Arakawa, T. and Timasheff, S.N. (1982) Stabilization of protein structure by sugars. Biochemistry, 21, 6536–6544.

52. Gekko, K. and Timasheff, S.N. (1981) Mechanism of protein stabilization by glycerol: preferential hydration in glycerol-water mixtures. Biochemistry, 20, 4667–4676.

53. Lee, L.L. and Lee, J.C. (1987) Thermal stability of proteins in the presence of poly(ethylene glycols). Biochemistry, 26, 7813–7819.

54. Arakawa, T. and Timasheff, S.N. (1985) The stabilization of proteins by osmolytes. Biophys J, 47, 411–414.

55. Timasheff, S.N. (1998) Control of protein stability and reactions by weakly interacting cosolvents: The simplicity of the complicated. Advances in Protein Chemistry, Vol 51, 51, 355–432.

56. Timasheff, S.N. (1993) The Control of Protein Stability and Association by Weak-Interactions with Water - How Do Solvents Affect These Processes. Annu Rev Bioph Biom, 22, 67–97.

57. Davis-Searles, P.R., Saunders, A.J., Erie, D.A., Winzor, D.J. and Pielak, G.J. (2001) Interpreting the effects of small uncharged solutes on protein-folding equilibria. Annu Rev Biophys Biomol Struct, 30, 271–306.

58. Saunders, A.J., Davis-Searles, P.R., Allen, D.L., Pielak, G.J. and Erie, D.A. (2000) Osmolyte-induced changes in protein conformational equilibria. Biopolymers, 53, 293–307.

59. Zhou, H.X., Rivas, G. and Minton, A.P. (2008) Macromolecular crowding and confinement: biochemical, biophysical, and potential physiological consequences. Annu Rev Biophys, 37, 375–397.

60. Knowles, D.B., LaCroix, A.S., Deines, N.F., Shkel, I. and Record, M.T., Jr. (2011) Separation of preferential interaction and excluded volume effects on DNA duplex and hairpin stability. Proc Natl Acad Sci U S A, 108, 12699–12704.

61. Knowles, D.B., Shkel, I.A., Phan, N.M., Sternke, M., Lingeman, E., Cheng, X., Cheng, L., O’Connor, K. and Record, M.T. (2015) Chemical Interactions of Polyethylene Glycols (PEGs) and Glycerol with Protein Functional Groups: Applications to Effects of PEG and Glycerol on Protein Processes. Biochemistry, 54, 3528–3542.

62. Bolen, D.W. and Baskakov, I.V. (2001) The osmophobic effect: natural selection of a thermodynamic force in protein folding. J Mol Biol, 310, 955–963.

63. Record, M.T., Jr., Guinn, E., Pegram, L. and Capp, M. (2013) Introductory lecture: interpreting and predicting Hofmeister salt ion and solute effects on biopolymer and model processes using the solute partitioning model. Faraday Discuss, 160, 9–44; discussion 103-120.

64. Bruck, I. and O’Donnell, M. (2001) The ring-type polymerase sliding clamp family. Genome Biology, 2.

65. Myers, J.K., Pace, C.N. and Scholtz, J.M. (1995) Denaturant M-Values and Heat-Capacity Changes - Relation to Changes in Accessible Surface-Areas of Protein Unfolding. Protein Sci, 4, 2138–2148.

66. Capp, M.W., Pegram, L.M., Saecker, R.M., Kratz, M., Riccardi, D., Wendorff, T., Cannon, J.G. and Record, M.T., Jr. (2009) Interactions of the osmolyte glycine betaine with molecular surfaces in water: thermodynamics, structural interpretation, and prediction of m-values. Biochemistry, 48, 10372–10379.

67. Jeruzalmi, D., Yurieva, O., Zhao, Y.X., Young, M., Stewart, J., Hingorani, M., O’Donnell, M. and Kuriyan, J. (2001) Mechanism of processivity clamp opening by the delta subunit wrench of the clamp loader complex of E-coli DNA polymerase III. Cell, 106, 417–428.

68. Burgers, P.M., Kornberg, A. and Sakakibara, Y. (1981) The dnaN gene codes for the beta subunit of DNA polymerase III holoenzyme of escherichia coli. Proc Natl Acad Sci U S A, 78, 5391–5395.

69. Leu, F.P., Hingorani, M.M., Turner, J. and O’Donnell, M. (2000) The delta subunit of DNA polymerase III holoenzyme serves as a sliding clamp unloader in Escherichia coli. J Biol Chem, 275, 34609–34618.

70. Kubitschek, H.E. and Friske, J.A. (1986) Determination of bacterial cell volume with the Coulter Counter. J Bacteriol, 168, 1466–1467.

71. Levin, P.A. and Angert, E.R. (2015) Small but Mighty: Cell Size and Bacteria. Cold Spring Harb Perspect Biol, 7, a019216.

