## Supplemental Information for "Potassium Glutamate and Glycine Betaine Induce Self-Assembly of Sliding Clamps into Higher Order Oligomers"

**Table of Contents**


| | |
|---|---|
| Figure S1: Apparent diffusion times measured in NaGlu vs KGlu | Page S2 |
| Figure S2: Control experiments with different dyes. β-I305C-TMR vs β-I305C-A546 | Pages S2-S3 |
| Figure S3: Control experiments with different labeling positions. β-I305C-TMR, β-S109C-TMR and β-C333-TMR. | Pages S3-S4 |
| Figure S4: Reversibility (high KGlu to low KGlu) | Page S5 |
| Calculation of $K_{\beta_2}$ | Pages S5-S6 |
| PyMOL modeling and Fig. S5 | Pages S6-S7 |

#### Supplemental Figure S1

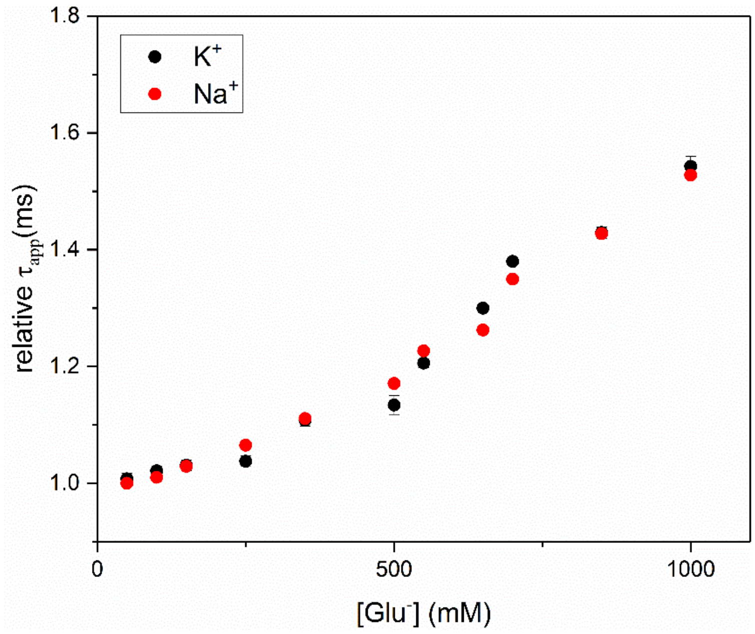


**Figure S1:** Apparent diffusion times normalized to the initial value measured in KGlu (reported in Fig. 2A) and in NaGlu.

#### Supplemental Figure S2

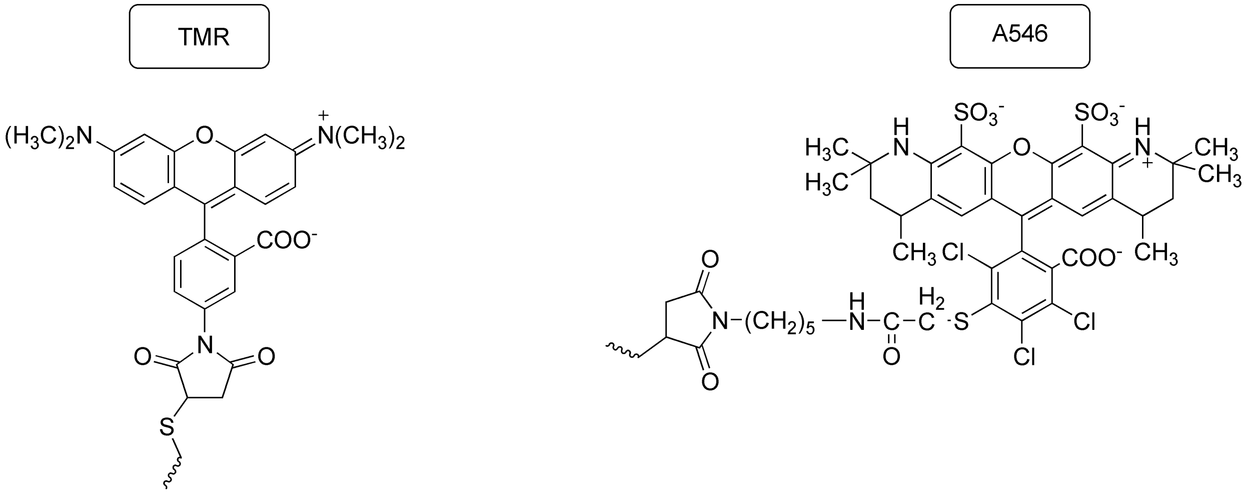

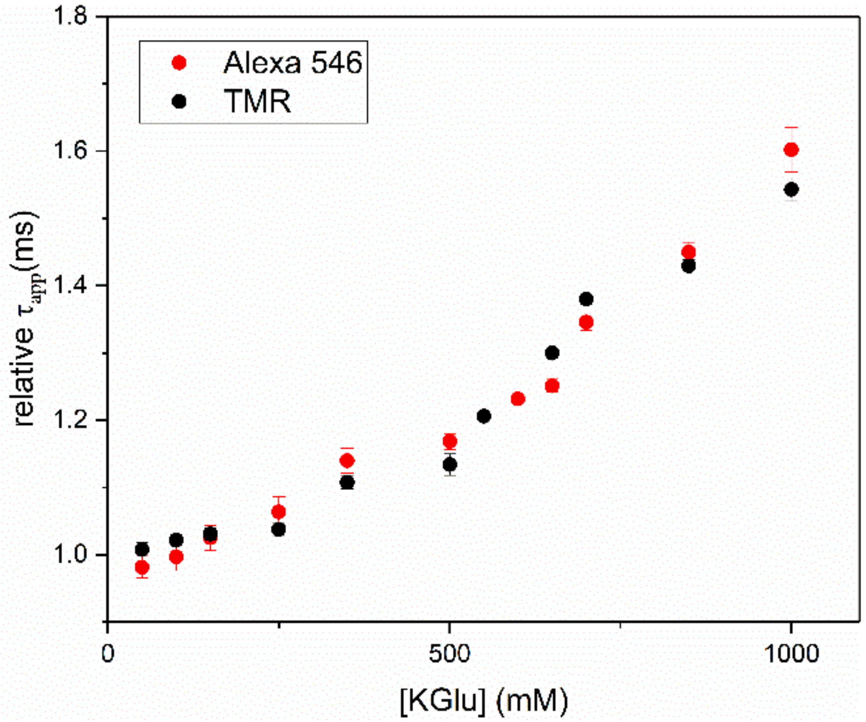


**Figure S2:** Apparent diffusion times normalized to the initial value measured with TMR- and Alexa 546-labeled β clamps (β-I305C-TMR and β-I305C-A546). The location of the fluorophore is the same in both cases (I305C).

#### Supplemental Figure S3

The same experiments reported in Fig. 2A were performed with clamps labeled with TMR in two other positions:

1) β-I305C-TMR (same data reported in Fig. 2A) contains the following mutations: C260S + C333S (background mutations) and I305C (for labeling with TMR-maleimide).

2) β-S109C-TMR contains the following mutations: C260S + C333S (background mutations) and S109C (for labeling with TMR-maleimide).

3) β-C333-TMR contains only one background mutation (C260A). The native amino acid Cys-333 was used for labeling with TMR-maleimide.

The amino acids labeled in each case are highlighted in the following figure (red = C333, blue=S109C, grey = I305C).

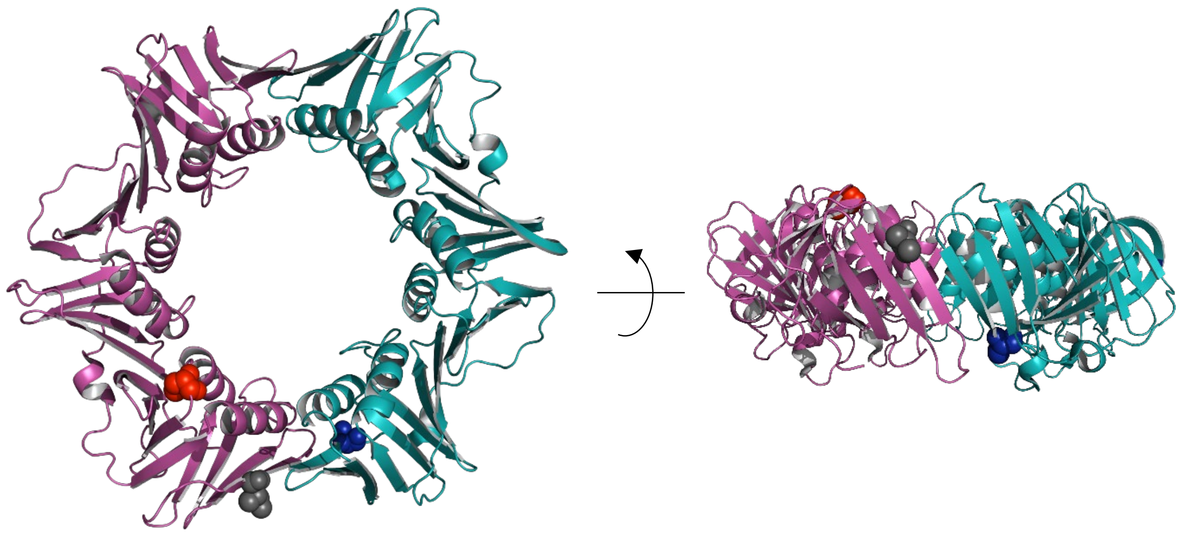

Relative diffusion times (normalized to the value measured without KGlu) are shown in the following Figure:

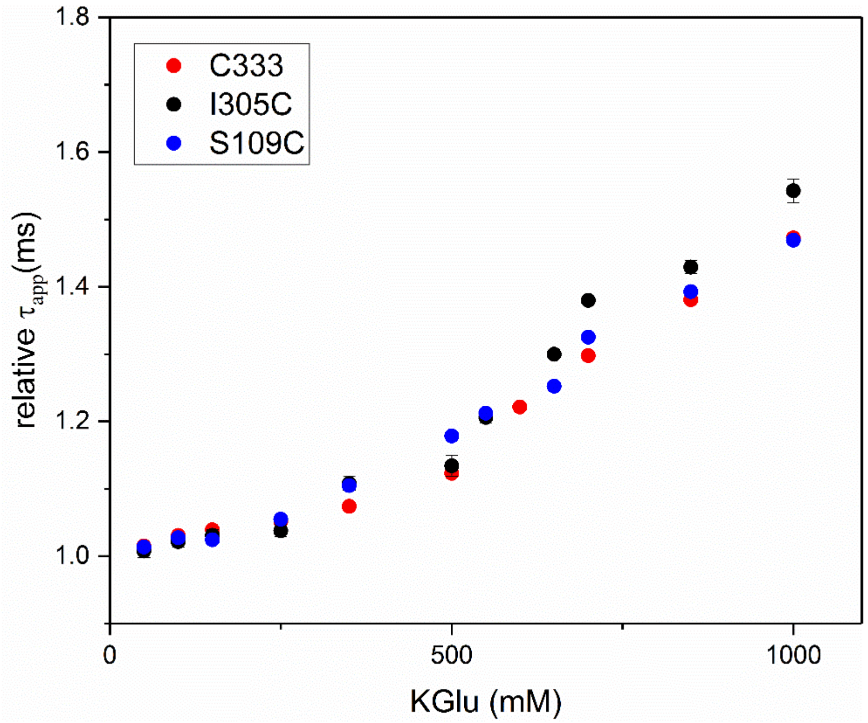

#### Supplemental Figure S4

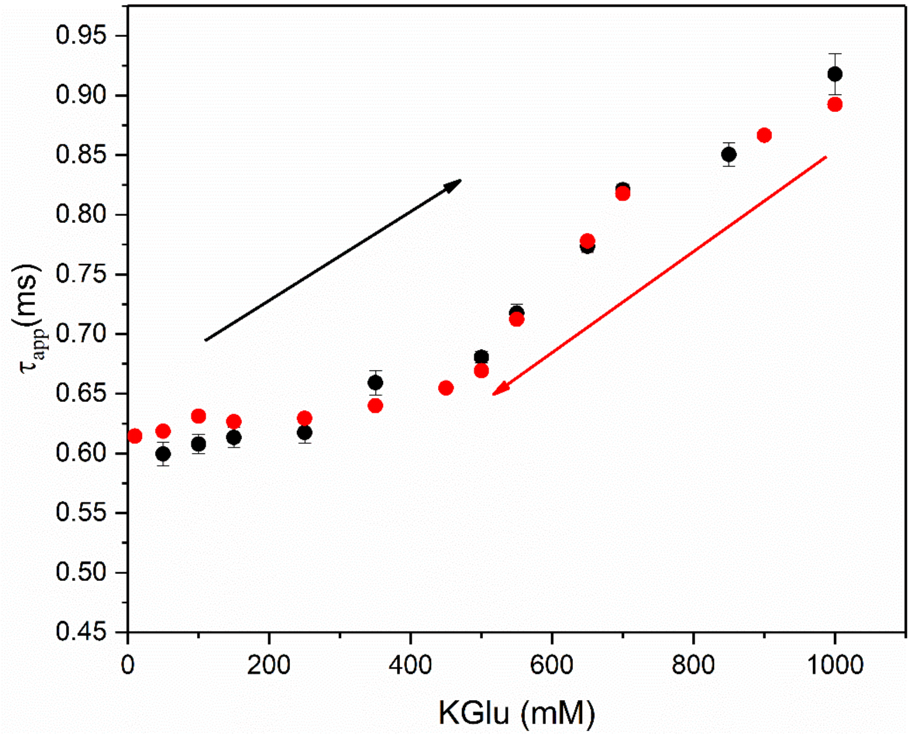


**Figure S4:** Apparent diffusion times measured by adding KGlu to β-I305C-TMR (black, same data presented in Fig. 2A) and by removing KGlu by dilution (red). For the experiments reported in black, the protein was diluted from a concentrated stock containing no KGlu into a buffer containing the desired concentration of Glu. For the experiments reported in red, the protein was initially dissolved in 1 M KGlu, and was then diluted to 1 nM using a buffer containing variable concentrations of KGlu to achieve the desired final Glu concentration.

#### Calculation of $K_{\beta_2}$

Here we use the formalism previously developed by Kanno and Levitus[1] to calculate the dissociation equilibrium constant of $\beta_2$ from the measured apparent diffusion times ($\tau_{app}$) at different KGlu concentrations. This section assumes that only monomers (i.e. clamps, β) and dimers (i.e. a stack of two clamps, $\beta_2$) exist in equilibrium. As discussed in the manuscript, this is only the case at moderate Glu concentrations.

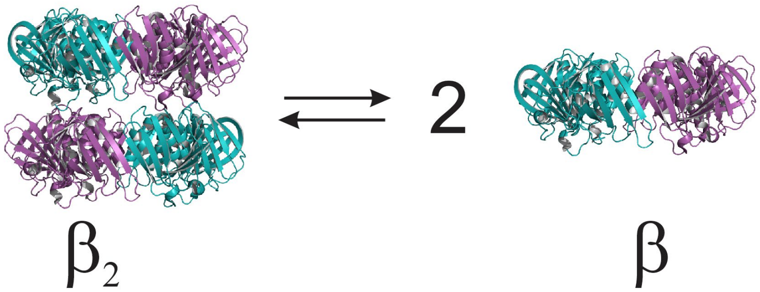


If the equilibrium between dimers (D) and monomers (M) is described by a single dissociation constant $K_d$, this constant can be expressed in terms of the degree of dissociation ($\alpha = C_M/C_0$) and the total protein concentration ($C_0$, expressed in terms of monomers):

$$K_d = \frac{2\alpha^2 C_0}{1-\alpha} \qquad \text{(S1)}$$

The measured apparent diffusion time ($\tau_{app}$) can be expressed as:[1]

$$\tau_{App}^2 - \left[1 - 2\frac{\alpha}{1+(1-\alpha)f}\right](\tau_2 - \tau_1)\tau_{App} - \tau_1\tau_2 = 0 \qquad \text{(S2)}$$

where $\tau_{1,2}$ are the diffusion times of the monomer and dimer, respectively, and $f$ is the labeling efficiency ($0 < f < 1$).

The degree of dissociation of 1 nM solutions of β were calculated at each KGlu concentration using Eq. S2 and the measured $\tau_{app}$ values of Fig. 2A. We stress that the term "degree of dissociation" refers to the equilibrium between $\beta_2$ and β, and not to the dissociation of the dimeric clamp itself, which we studied in previous work. As discussed in the manuscript, clamp dissociation was determined to be negligible in the timescales of the experiments reported in this manuscript. The value of $\tau_1$ in Eq. S2 was set to 0.598 ms, as measured in the experiment with 1 nM β in 50 mM NaCl/TRIS buffer. The value of $\tau_2$ was set to $1.208\tau_1$ based on estimates of the diffusion coefficient of β and $\beta_2$ using the computer program HYDROPRO.[2] The pdb file of $\beta_2$ (shown in Fig. S5) was created in PyMOL. Finally, $K_d$ was calculated from α using Eq. S1 ($C_0$ = 1nM), and association constants ($K_{\beta_2}$) were calculated as 1/ $K_d$.

### PyMOL modeling

Pdb structures of model dimers of β were created in PyMOL from two identical copies of β (pdb id: 1MMI) using the following script:

```
fetch 1mmi
copy obj01, 1mmi
cealign 1mmi, obj01
orient
rotate x, 180, 1mmi
translate [0,0,d], 1mmi
rotate z,70, 1mmi
alter (obj01 and chain A), chain='C'
```

```
alter (obj01 and chain B), chain='D'
select all1, all
create dimer,all1
save filename.pdb, dimer
```

where d is the displacement between the two rings (32Å, 33Å, or 34Å)

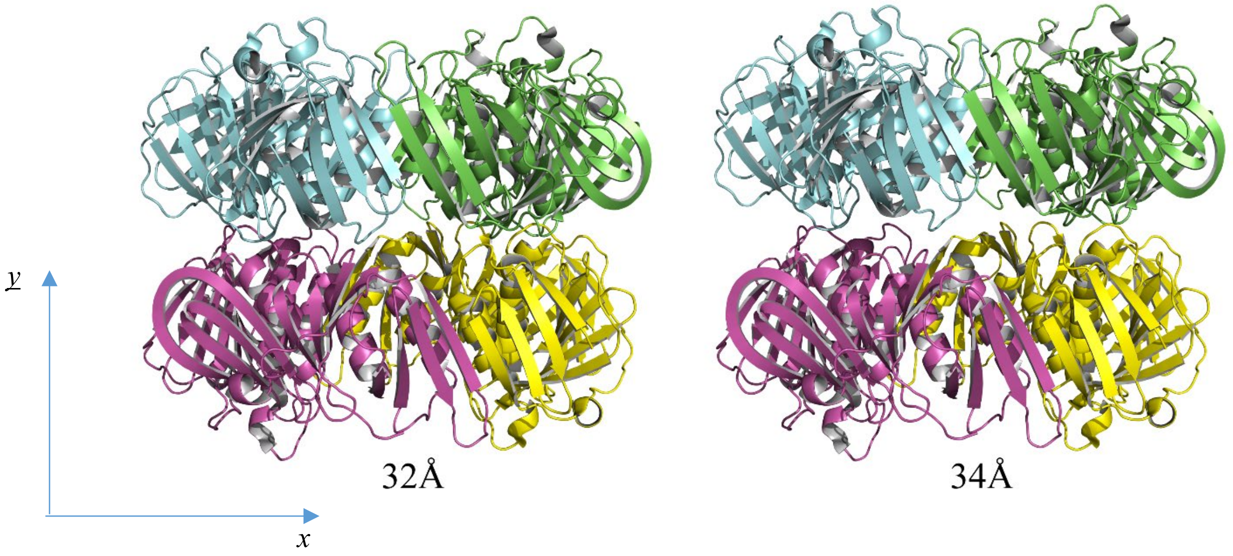


**Figure S5:** Pymol models of f $\beta_2$. The distances refer to the displacements d defined in the code above.


(1) Kanno, D. M.; Levitus, M. Protein Oligomerization Equilibria and Kinetics Investigated by Fluorescence Correlation Spectroscopy: A Mathematical Treatment. *J Phys Chem B* **2014**, *118*, 12404-12415.

(2) Ortega, A.; Amoros, D.; Garcia de la Torre, J. Prediction of hydrodynamic and other solution properties of rigid proteins from atomic- and residue-level models. *Biophys J* **2011**, *101*, 892-898.
